# Fibernet 2.0: An Automatic Neural Network Based Tool for Clustering White Matter Fibers in the Brain

**DOI:** 10.1101/210781

**Authors:** Vikash Gupta, Sophia I. Thomopoulos, Conor K. Corbin, Faisal Rashid, Paul M. Thompson

## Abstract

The brain’s white matter fiber tracts are impaired in a range of common and devastating conditions, from Alzheimer’s disease to brain trauma, and in developmental disorders such as autism and neurogenetic syndromes. Many studies now examine the connectivity and microstructure of the brain’s neural pathways, spurring the development of algorithms to extract and measure tracts and fiber bundles. Clustering white matter (WM) fibers, from whole-brain tractography, into anatomically meaningful bundles is still a challenging problem. Existing tract segmentation methods use atlases or regions of interest (ROI) or unsupervised spectral clustering. Even so, atlas-based segmentation does not always partition the brain into a set of recognizable fiber bundles. Deep learning techniques can be applied to automatically segment and cluster white matter fibers. Here we propose a robust approach using convolutional neural networks (CNNs) to learn shape features of the fiber bundles, which we then exploit to cluster WM fibers into bundles. In a range of tests across diverse fiber bundles, we illustrate the accuracy of our method, and its ability to suppress false positive fibers.

## 1. INTRODUCTION

Diffusion-weighted MRI offers a non-invasive method to study the micro-structure of the brain; when processed using whole-brain tractography methods, patterns of anatomical connections can be mapped between multiple brain regions, and these patterns can be analyzed to understand disease effects. Tracts, or axonal bundles in the white matter (WM), are neural pathways that interconnect different parts of the brain. Diffusion tensor imaging and its higher-order extensions (such as q-space imaging, or diffusion spectrum imaging), are used increasingly to study WM pathways. To study disease effects on specific regions within the white matter, it can be useful to cluster the fibers into anatomically meaningful fiber bundles. Fiber clustering offers a first step towards statistics on large datasets, where it may not be feasible to segment the WM fibers by hand for every single subject. Some widely used methods to partition the WM fibers are based on region of interest (ROI) segmentation. ROIs are delineated, automatically or by hand, on T1-weighted anatomical images, which are co-registered with the fractional anisotropy (FA) images. These ROIs are then used to “seed” or filter sets of streamlines from tractography, isolating subsets of fibers for subsequent analyses. Even when this is done, it is still hard to define the major fiber pathways of the brain based on any one set of ROIs; individual pathways may also merge, branch and intersect, making them hard to partition correctly using a deformable atlas or template. Basing tract definitions on the ROIs they intersect can lead to incorrect splitting or fusion of fiber bundles, or omission of some fibers that belong to the same bundle. The fiber clustering problem is extremely challenging, and has been tackled for over a decade using unsupervised methods such as spectral clustering [1, 2] and more recently using model-based supervised methods [3]. Inspired by the success of convolutional neural networks in the computer vision community [4], recently CNNs have begun to be tested for clustering white matter fibers in the brain. In this paper, we extend work we presented in [5], to better handle false positive fibers. A whole brain tractography algorithm produces as much as four times the false-positive fibers (pathways that are incorrectly tracked and do not correspond to true anatomy) when compared to true fibers [6]. A variety of factors can promote the generation of these spurious fibers: low signal-to-noise (SNR), low spatial, spectral, or angular resolution in the diffusion images, or an abrupt drop in fractional anisotropy (FA) values (and uncertainty in fitting dominant fiber directions) at fiber crossings, among other factors. To this end, false positive fibers cause serious problems for fiber clustering methods, and for any downstream analyses based on tractography. In our prior work [5], we were not able to compensate for spurious fibers, but we presented a proof of concept that CNN based neural network architectures might be used for fiber clustering tasks. In this paper, we build a more practical and robust approach to reduce the effects of false positive streamlines.

## 2. VOLUMETRIC PARAMETERIZATION

To design a neural network to learn features for classifying WM fibers, it is important to impose a consistent set of co-ordinates across subjects. Unlike positions in a 3D voxel lattice, fiber paths in the brain are geometric entities and depend on the geometric coordinate system used for the image acquisition (such as the image direction cosine matrix, the image origin, and voxel resolution). These parameters vary across different datasets and scanners. To resolve this, we use a volumetric parameterization technique, proposed in [5]. This parameterization may also be thought of as a normalization process for the fiber tracts. The method uses a 3D harmonic function to generate level-sets based on a chosen *shapecenter*, which is designed to be anatomically consistent across subjects. Streamlines orthogonal to the level-sets are computed by solving an ordinary differential equation. The resulting nonlinear grid (figure 1) is used for parameterizing all the fiber tracts to this new coordinate system. The curvilinear coordinates also provide a convenient spherical domain for embedding the brain volume, prior to learning statistical distributions governing tract variability across subjects.

**Fig. 1.**
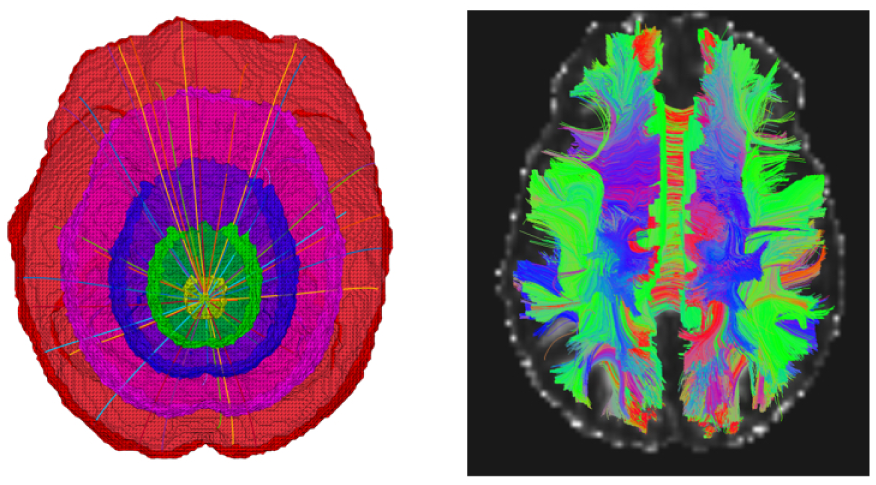
**Left:** Volumetric parameterization with level sets and streamlines. **Right:** Whole brain tractography.

## 3. DATA AND METHODS

The data analyzed in this study is a publicly available dataset from the Parkinson’s Progression Markers Initiative (PPMI - www.ppmi-info.org). For each subject, a high resolution T1-weighted anatomical image was acquired along with a set of diffusion-weighted images (DWI) using a 3 Tesla Siemens scanner. The T1-weighted scan consists of 256 × 240 × 176 voxels with an isotropic resolution of 1 mm. The DWIs were also acquired using the same scanner and consisted of 116 × 116 × 72 voxels with an isotropic resolution of 2 mm. For the diffusion data, 64 images were acquired along 64 gradient directions (b-value = 1000 *s/mm*^2^) and one T2-weighted B0 image was acquired with no diffusion sensitization.

### 3.1. Data Preprocessing

Each subject’s diffusion weighted images (DWIs) were corrected for head motion and eddy current distortions, using FSL tools. Intra-subject linear registration was used to align the undiffused B0 and the anatomical T1w images. The T1w images were aligned to the MNI template using an affine registration. All the registrations were performed using FSL’s *FLIRT* utility. For computing the volumetric parameterization described in section 2, a common anatomically equivalent point, called *shapecenter* should be identified in all the subject space. This point is identified as (98, 113, 111) on the MNI template. For each subject, the transformations computed in the previous step are combined in order and inverted. The inverse transformation was used to locate the shapecenter in each subject’s diffusion space. Based on this shapecenter, the volumetric parameterization was computed.

### 3.2. Tractography and Data Augmentation

Camino’s fiber reconstruction algorithm called the Probabilistic Index of Connectivity (PICo) [7] was used for fiber reconstruction. Seed points were chosen at voxels with an FA value greater than 0.2. Monte Carlo simulation was used to generate streamlines proceeding from the seed points throughout the entire brain [8]. Streamline fiber tracking followed the voxel-wise probability distribution function profile with the Euler interpolation method, for 7 iterations at each seed point. The maximum fiber turning angle was set to 60°/voxel, and tracing stopped at any voxel whose FA was less than 0.2.

The tractography results were first filtered based on ROI definitions available from FreeSurfer segmentations. Given the domain expertise of our neuroanatomist, we are aware of fiber bundles that interconnect or pass through any given set of ROIs. Union and intersection operations are performed on the fiber bundles in conjunction with the ROIs to perform a coarse filtering of the fiber tracts. However, as many spurious fibers are present, such an operation still leaves a considerable number of false-positive fiber tracts in the datasets.

To train our neural network, we selected four subjects at random from the PPMI database. After tractography and coarse fiber segmentation, false positive fibers were manually removed by a neuroanatomy expert. Thus, for each set of fiber bundles we have *positive* and *negative* examples. This process also leads to a class imbalance issue for some of the fiber bundles - as shown in table 1. This issue runs both ways, i.e., for some fiber bundles the number of positive examples is greater than the negative ones and *vice versa*. If negative examples are fewer than the positive ones, the minority class is oversampled to match the majority class. On the other hand, if there are more negative examples, a small amount of “jitter” is added into the fiber tracts. To do this, the fiber tracts are convolved with 3 one-dimensional Gaussian filters along the three coordinate axes. Because of the additive property of Gaussian filters, this operation is equivalent to performing a convolution along the tract. In addition, the order of points on each fiber tract is flipped, and the flipped fibers are added to the set of streamlines available for training. Finally, these fiber tracts are parameterized using the method described in section 2. The parameterized tracts are used as feature vectors to train the neural network.

**Table. 1.** The table shows number of positive and negative examples available from four training subjects for each of the fiber bundles. The two rightmost columns show the true positive rate (TPR) and true negative rate (TNR) on the validation set. **Acronyms:** ATR: anterior thalamic radiations, CC: corpus callosum, PRCG: pre-central gyrus, POCG: post- central gyrus, IFO: inferior longitudinal fasciculus, SLF: superior longitudinal fasciculus, L/R: left/right

| Fiber Bundle | # +ve | # -ve | TPR | TNR |
| --- | --- | --- | --- | --- |
| ATR.L | 20748 | 3961 | 0.99 | 0.99 |
| CC.POCG | 18020 | 9714 | 0.99 | 0.99 |
| CC.TEMPORAL | 11258 | 11600 | 0.96 | 0.99 |
| CC.PARIETAL | 35106 | 14853 | 0.97 | 0.98 |
| CC.OCCIPITAL | 26568 | 14922 | 0.99 | 0.98 |
| CC.FRONTAL | 120817 | 39783 | 0.97 | 0.98 |
| CC.PRCG | 25702 | 9699 | 0.99 | 0.99 |
| IFO.L | 18429 | 5877 | 0.95 | 0.80 |
| IFO.R | 12920 | 2371 | 0.99 | 0.86 |
| SLF.L | 17826 | 5518 | 0.97 | 0.97 |

**Fig. 2.**
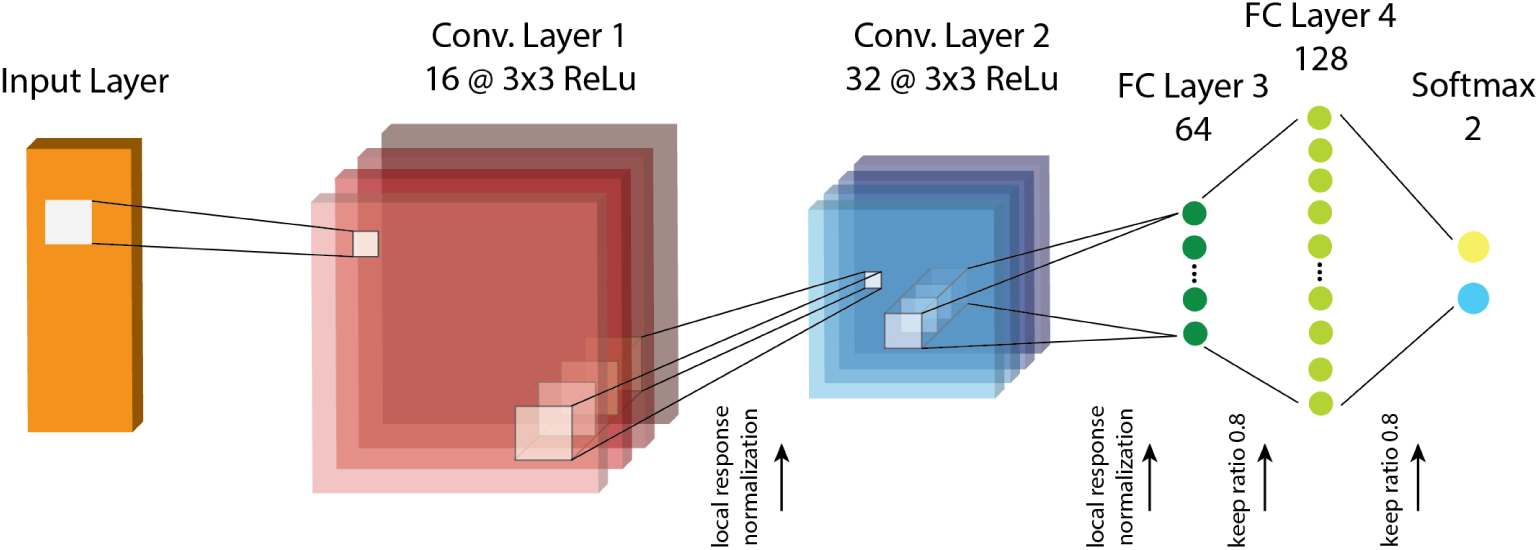
FiberNet 2.0: The neural network architecture.

## 4. FIBERNET 2.0

In this paper, we extended the workflow described in a previous version of FiberNET [5] for classifying WM fibers. These changes help us counter the presence of spurious fibers, which have been said to comprise a majority of fiber tracts computed using typical tractography algorithms [6]. As we are working with less input data and binary classification, we reduced the number of convolutional feature maps and the fully connected layers by half. In this version of FiberNET, the network contains 2 layers of 16 and 32 feature maps followed by 2 fully connected layers with 64 and 128 nodes. The last layer consists of a“softmax” layer with two classes. To prevent over-fitting, we use 80% keep-ratio, which randomly switches off neuronal units in each layer thereby reducing their influence at any particular iteration during back-propagation. For inducing nonlinearity, the activation function used in the convolutional layers and the fully connected layers are rectified linear units (ReLU) and hyperbolic tangent (tanh) respectively. A batch size of 3000 fiber tracts and a learning rate of 0.0001 was used for the optimization process. In the present work, deep learning was implemented using the TensorFlow version r0.11. We also employed a bootstrap aggregation for further generalization. Essentially the data is sampled with replacement a number of times (in this case 20) and a subset of the data is created. For each sampling, a neural network is trained with the above mentioned parameters. For a particular fiber a class probability is computed from each of the 20 models and the class label is assigned based on majority voting.

## 5. RESULTS

We trained our network for classifying ten tracts listed in table 1, In Figure 3, we show the results of classification for these different tracts. In the left column, one can see the result of ROI based segmentation and the right column shows the result of binary classification on one of subjects. The last two columns of table 1 show the true positive rate (sensitivity) and true negative rate (specificity) for the classification. We can see that, except for the inferior fronto-occipital fasciculus (IFO) the accuracy is fairly above 97%. The lower accuracy in the IFO could result from a class imbalance issue and is the subject of further investigation. We show the final classified fiber bundles in figure 3.

**Fig. 3.**
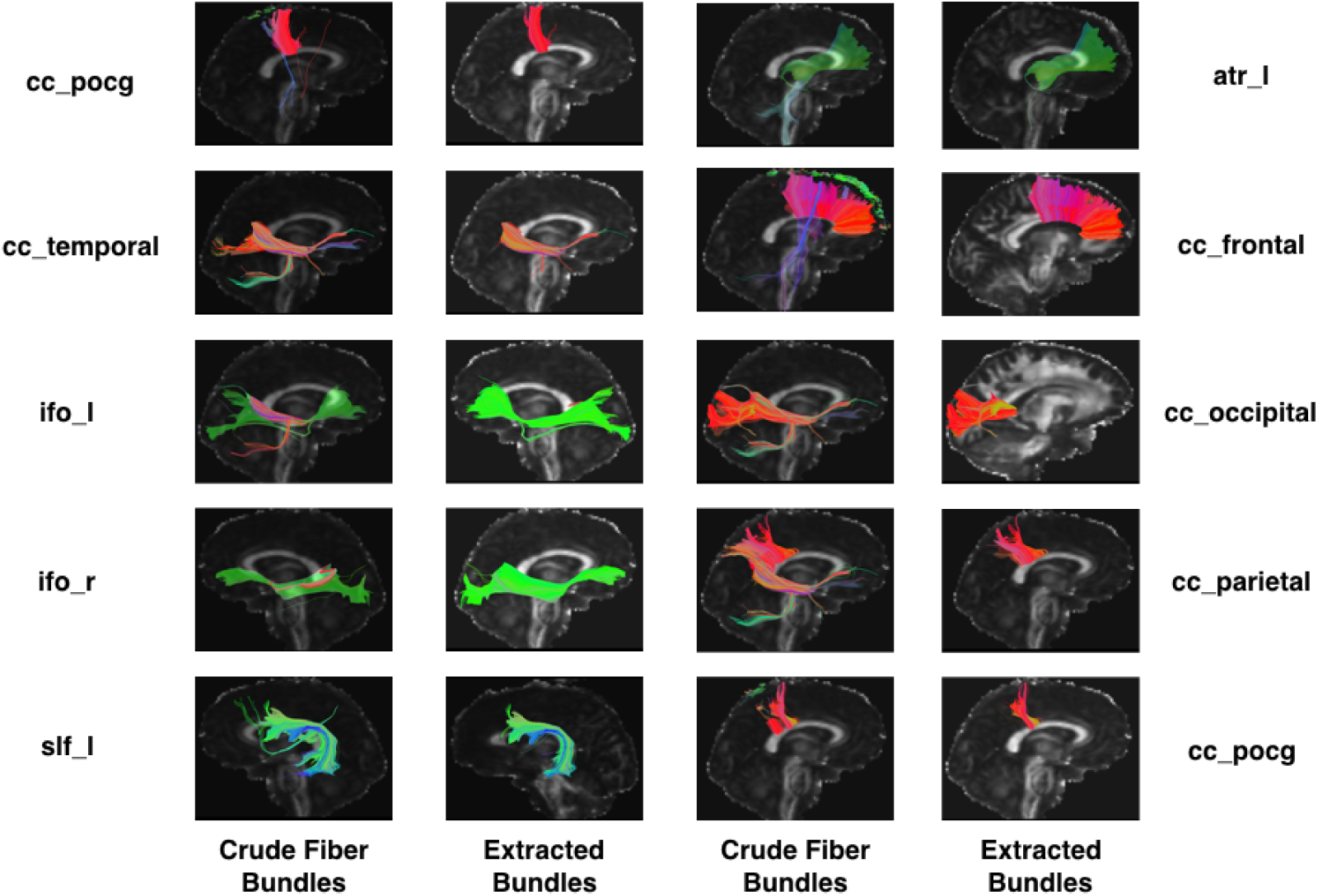
Results of binary classification on ten tracts for one particular subject chosen at random from the PPMI dataset

## 6. CONTRIBUTIONS AND CONCLUSIONS

In this paper, we presented a workflow based on our previous method FiberNET. The proposed workflow aims to remove false positive fibers from a fiber bundle segmented using ROI based segmentation tools. It is important to note that false-positive tracts dominate whole brain tractography results [6]. In [6], the authors show that even the different tractography algorithms are able to produce 90% of ground truth fibers, they produce nearly four times as much false positive fibers. Removing these false positives is absolutely critical to conducting a large-scale statistical study. The method we presented is designed to efficiently handle noisy whole-brain tractography data and yield anatomically relevant fiber tract bundles with high accuracy. The method is fully automated and is applied to the PPMI dataset, consisting of data from 226 subjects, and it produced promising results showing the generalizability of the method. In the future, we would like to test our model on multi-shell tractography data and test whether a model trained on a multi-shell data can be used to predict or cluster single shell tractography results.

